# Mapping Protein Numbers in Living Cells

**DOI:** 10.1101/2021.07.02.450806

**Authors:** Derek J Thirstrup, Jie Yao, Jamie Sherman, Irina A Mueller, Winfried Wiegraebe

**Affiliations:** Allen Institute for Cell Science, Seattle, WA, USA; Nautilus Biotechnologies, Seattle, WA, USA; Bruker Nano Surfaces & Metrology, Madison, WI, USA

## Abstract

We introduce a new, robust method to map the numbers of proteins in living cells. The method can be applied to laser scanning, spinning disk, and lattice light-sheet microscopes in a robust, reproducible, and scalable fashion. The method uses calibrated EGFP solutions that are imaged with the appropriate microscope modality to create a calibration curve that is then applied to convert the fluorescence intensities from 3D microscope images into molecule numbers. We applied this method to human induced pluripotent stem cells in which proteins representing key cellular structures were endogenously tagged with mEGFP. We used the ratio of mEGFP-tagged proteins to total proteins to create 3D maps of live cells showing the density of total proteins measured in molecules per µm^3^. The method opens the door to new quantitative single cell analyses of cellular protein numbers in the context of single cell gene expression, associations with cellular complexes, and changes in cellular behaviors. The method is capable of quantifying protein numbers, over three orders of magnitude, in the cytoplasm or within various cellular structures while offering the unique advantages of each microscopy modality.

## Introduction

The quantification of mRNA expression in whole cells, using single cell RNA-seq, and in regions of cells, using RNA FISH, is transforming our understanding of gene expression and how it determines cellular phenotypes. Analogous methodologies for measuring protein expression levels could be similarly transformative. They not only reveal the relations between gene and protein expression but also help to determine equilibria between soluble and bound molecules, reveal biological consequences of variations in concentration within cells and cell populations, and identify and understand changes in cellular behaviors.

In principle, the fluorescence intensities, measured by microscopes, can report the number of fluorescent protein (FP) tagged proteins. However, attempts to make these kinds of measurements are infrequent, used for a specific structure, or limited to a specialized approach and have not scaled into a general, useful tool for routine or high throughput studies [1]. Some examples include: 1) Stepwise photobleaching using total internal reflection fluorescence (TIRF) microscopy was used to quantify low abundance proteins in single cells [2, 3]. 2) Ratiometric comparisons using spinning disk confocal microscopy with internal or external standards were used to quantify protein numbers in centromeres and kinetochores in *Schizosaccharomyces pombe* and *Saccharomyces cerevisiae* using GFP-tagged viral particles or GFP-MotB protein in *Escherichia coli* [4, 5]. 3) Protein number in cell populations was estimated using quantitative immunoblots of purified fluorescent proteins and protein extracts of yeast strains with homozygous tagged target genes [6]. 4) Finally, the concentrations of EGFP-tagged proteins were estimated using fluorescence correlation spectroscopy (FCS) and related fluctuation methods to reveal oscillations in the amount of Cse4p, a centromeric histone protein, in *S. cerevisiae* during the cell cycle [7] and measure local concentrations of GFP-tagged proteins in cells [8-11].

These successes reveal the importance and utility of quantifying protein numbers in and within single cells, However, they also show the need for a more general approach applicable to higher eukaryotic cells that can be employed with commonly used imaging modalities and produce large scale data output. Previous studies that image serially diluted EGFP solutions and EGFP-expressing cells [12] point to such a method for mapping protein concentrations in mammalian cells.

We extended and developed this method taking advantage of the gene-edited human induced pluripotent stem cell (hiPSC) lines developed at the Allen Institute for Cell Science. In these cell lines, proteins representing major cellular structures were tagged endogenously on one allele of a single locus with a fluorescent protein [13]. Thus, fluorescence intensities captured in a microscope image were a direct readout of the absolute number of proteins expressed from that genomic locus, providing an estimate of protein concentrations at the endogenous level. We optimized the method to be robust and applicable in 3D, in a high-throughput setting and across different microscope modalities, including enhanced resolution laser scanning confocal (LSM), spinning disk (SD) confocal, and lattice light-sheet (LLS) microscopy. In doing this, we also developed methods of correcting systematic intensity differences across imaging fields for SD and LLS microscopes and corrected for differential expression of the tagged and untagged alleles.

The procedure consists of five steps (Figure 1): 1) Measure the concentrations of EGFP in a dilution series typical for the proteins of interest via absorption spectroscopy, immunoblotting or FCS. 2) Optimize settings on each microscope modality and acquire cell images. 3) Image the EGFP dilution series using the same settings, creating a calibration curve relating fluorescence intensities to fluorophore concentrations. On SD and LLS microscopes, we accounted for uneven illumination profiles using per-pixel calibration. 4) Apply the calibration curve to the cell images. 5) Correct for the ratio between mEGFP-tagged and total protein expression when using monoallelic tagging. This workflow gave similar results across several microscope platforms. We applied this method to 15 different hiPSC lines with different cellular structures labeled with mEGFP. We found protein densities ranging from less than 50 molecules/µm^3^ (peroxisomal protein PMP34) to 45,000 molecules/µm^3^ (nucleophosmin, see Figure 3), indicating that this method can measure local densities of protein numbers inside living cells over three orders of magnitude.

**Figure 1.**
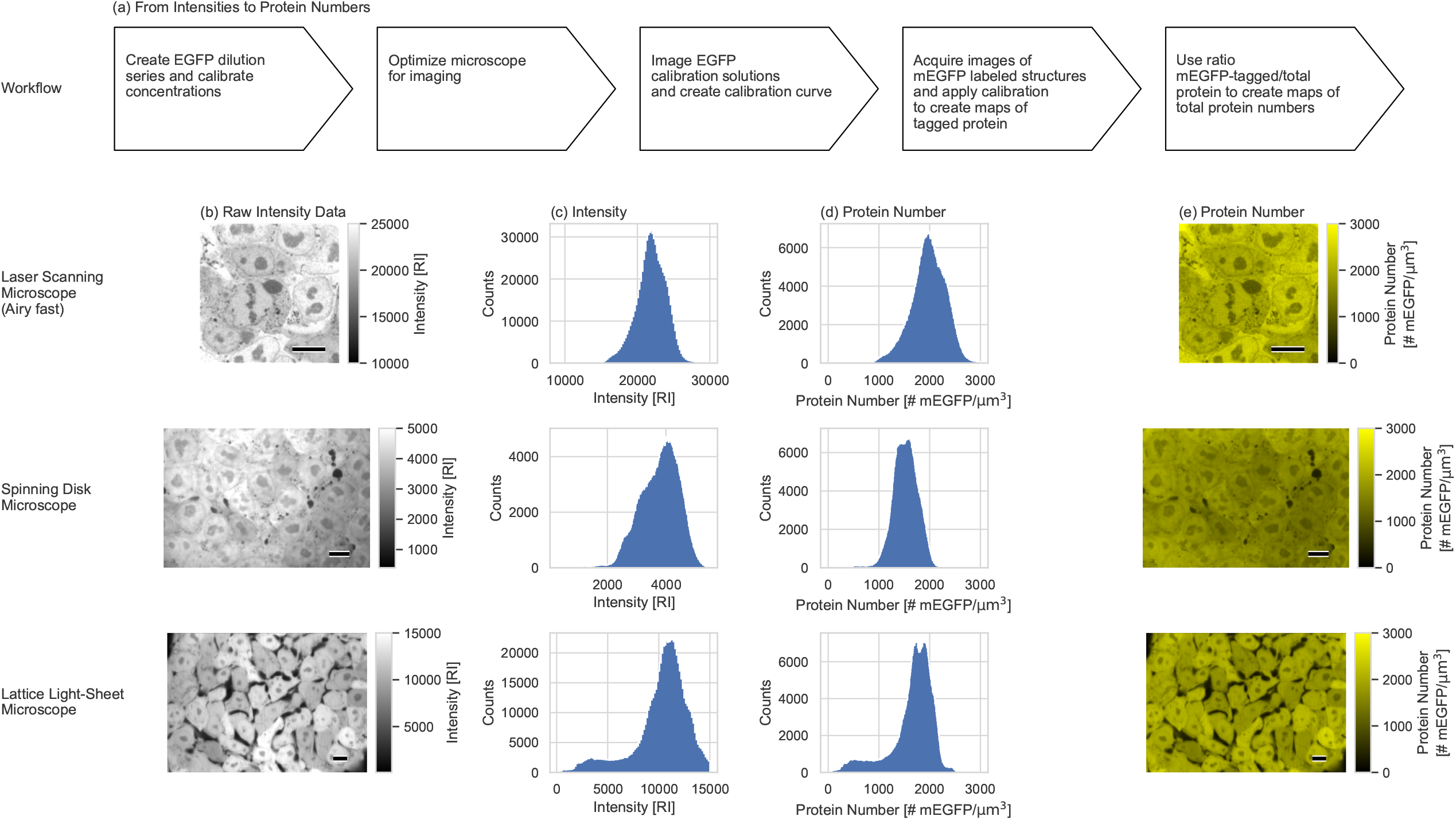
Calibration of fluorescence images. **(a)** Workflow for the calibration of microscope images. **(b)** Representative, single z-slice fluorescence images, from 3D images of live hiPSCs acquired with laser scanning microscope (LSM) with Airy fast detector (top), spinning disk microscope (SD, middle), and lattice light-sheet microscope (LLS, bottom). The cells expressed mEGFP as a free cytosolic protein driven by a CAGGS promoter from the AAVS1 safe harbor locus. Images were acquired with different microscopes and cells resulting in expression level and acquisition variation. The scale bars are 10 μm. The grey level bars were scaled in relative intensity units [RI]. The shading at the edges of the SD images caused by the Gaussian illumination profile and the detector specific offset are typical for SD microscopes. **(c)** Histograms of pixel intensities in relative intensity units [RI] for 3D image slices are shown in (b). The different laser powers, detector offsets and gain settings, illumination profiles and optical characteristics produced differences in offset and width of the intensity distributions for the different microscopes. **(d)** Protein number histograms of cytosolic mEGFP for image slices are shown in (b). All data are expressed as the number of proteins (mEGFP molecules) per μm^3^. The protein number distributions were very similar, independent of the microscope type used. **(e)** Images based on (b) showing protein numbers for cytosolic mEGFP. The color bars were scaled in number of mEGFP molecules per μm^3^. Artifacts from uneven illumination, especially visible in the spinning disk image in (b), were corrected, see Methods. The scale bars are 10 μm.

## Results and Discussion

### Calibration of microscope images

To calibrate the intensities from our platform, we first prepared EGFP solutions in the range of 10 nM to 2 μM, as these are the anticipated concentrations of most proteins in cells. We quantified the concentrations using absorption at 280 nm and 488 nm. We also measured EGFP concentrations using FCS [14, 15] (Supplemental Figure S2, also see Methods). The 280 nm and 488 nm absorption agreed to within about 20%; this difference is not surprising since the former estimates total protein and the latter estimates active fluorophore concentration. The concentration values measured from FCS were in between the 280 nm and 488nm concentrations (data not shown) and resulted in good linear correlation with the nominal EGFP concentrations (Supplemental Figure S2c). From multiple days of experiments performed over a year, slopes of the linear fit between FCS concentrations and nominal EGFP concentrations were always between 0.8 to 1.2 (mean ± SD: 1.03 ± 0.11). Therefore, FCS is capable of directly measuring EGFP concentrations at each microwell we used for calibration imaging with an uncertainty within 20%. We used the FCS measurement for our studies because it could be applied daily in our high throughput microscopy pipeline. Nonetheless, this calibration method can be generalized by using EGFP stock concentration values derived from other methods such as UV-VIS absorption spectroscopy if an FCS capable microscope is not available (see Methods for discussion).

To establish the FCS based image calibration method, we used an LSM because of its superior resolution and image quality (Figure 1b). It was equipped with an Airy fast detector to increase spatial resolution and imaging speed. We imaged Alexa 488 dye solutions and found no intensity variations across the field-of-view (FOV) (data not shown). Thus, we concluded that we could use identical calibration parameters for all pixels in the image. Then, we varied the concentrations of Alexa 488 and imaged the solutions with different exposure times. In both cases the detector response was linear (Supplemental Figure S1). Based on these findings, we imaged EGFP solutions of known concentrations using parameters optimized for cell imaging, fitting a linear regression to the intensity data and thus creating a calibration curve (Figure 2b), which we then applied to cell images of mEGFP-tagged proteins in hiPSCs. This quantitative method significantly facilitates the conversion of raw microscope images to calibrated spatial maps of protein number per μm^3^, providing absolute rather than relative protein densities in living cells.

**Figure 2.**
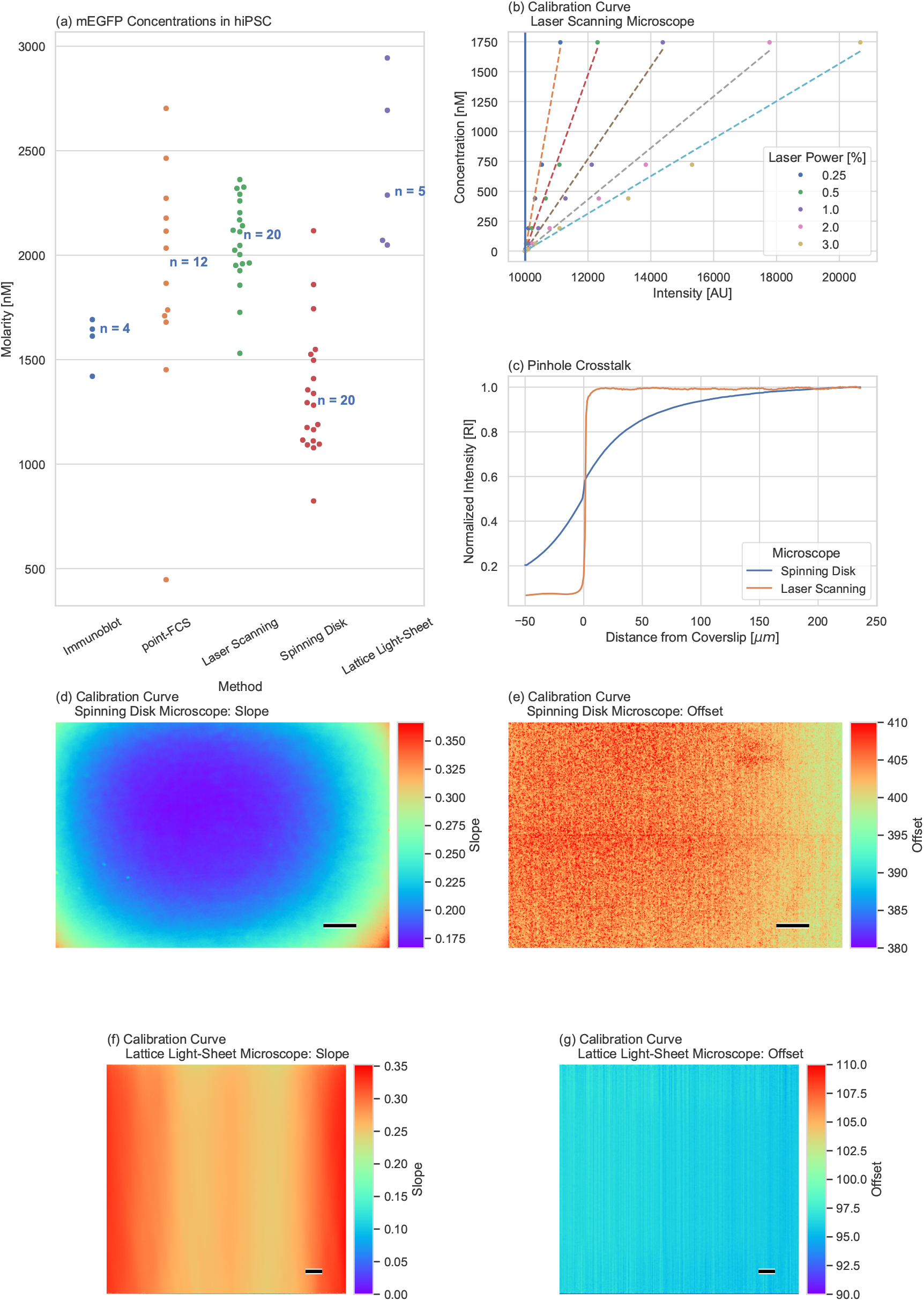
Calibration of image intensities to protein (fluorophore) number. **(a)** Average mEGFP concentrations of the same cell line shown in Figure 1 determined from (from left to right) quantitative immunoblots, point-FCS in single cells, and the protein number calibration method applied to LSM, SD, and LLS microscopy images. Each dot represents one cell pellet replicate (immunoblot), one cell (point-FCS) or one field-of-view (FOV) (microscopy methods). Each LSM image contains an average of three cells. **(b)** Calibration curves for LSM acquired at different laser powers. The same calibration curve was applied to each pixel in an xyz image stack. The dotted lines represent the slope of the calibration, the solid vertical line is the offset. **(c)** Intensity measured with SD microscope and LSM at different z-distances away from the coverslip (solution/glass interface). Images were acquired for Alexa 488 solutions of different concentrations and locations above the coverslip, and the intensities were averaged and normalized over the whole FOV. **(d, e)** Calibration curve for SD microscope. Slope (d) and offset (e) depended on the position within the FOV (xy plane). Scale bars are 10 μm. **(f, g)** Calibration curve for LLS microscope. Slope (f) and offset (g) depended on the position within the FOV (xy plane). Scale bars are 10 μm.

The lower detection limit of the method is determined by detector noise, scattering and autofluorescence. The nonuniform autofluorescence in cells presented a challenge (Figure 3o); it varied from the equivalent of 1.2 ± 0.6 proteins/μm^3^ in the background of a typical FOV, to peak values of 13 ± 3 proteins/μm^3^ with an average of 2 ± 2 proteins/μm^3^ for the whole FOV of wild type iPS cells. To determine the limits of detection, we used the well established level-of-detection (LOD), which defines the threshold that separates signal from background noise [16-18]. In general, the LOD was equivalent to 19 proteins per μm^3^, which is higher than the peak autofluorescence values we typically saw in hiPSC (Figure 3o). Using a non-clonal hiPSC line expressing low level mEGFP, we could clearly identify mEGFP-expressing cells in which cytosolic mEGFP density ranges between 25 and 100 proteins/µm^3^ (Figure 3n). Thus, we assessed that the practical detection limit of mEGFP in cells in our method is lower than 25 proteins/µm^3^.

**Figure 3.**
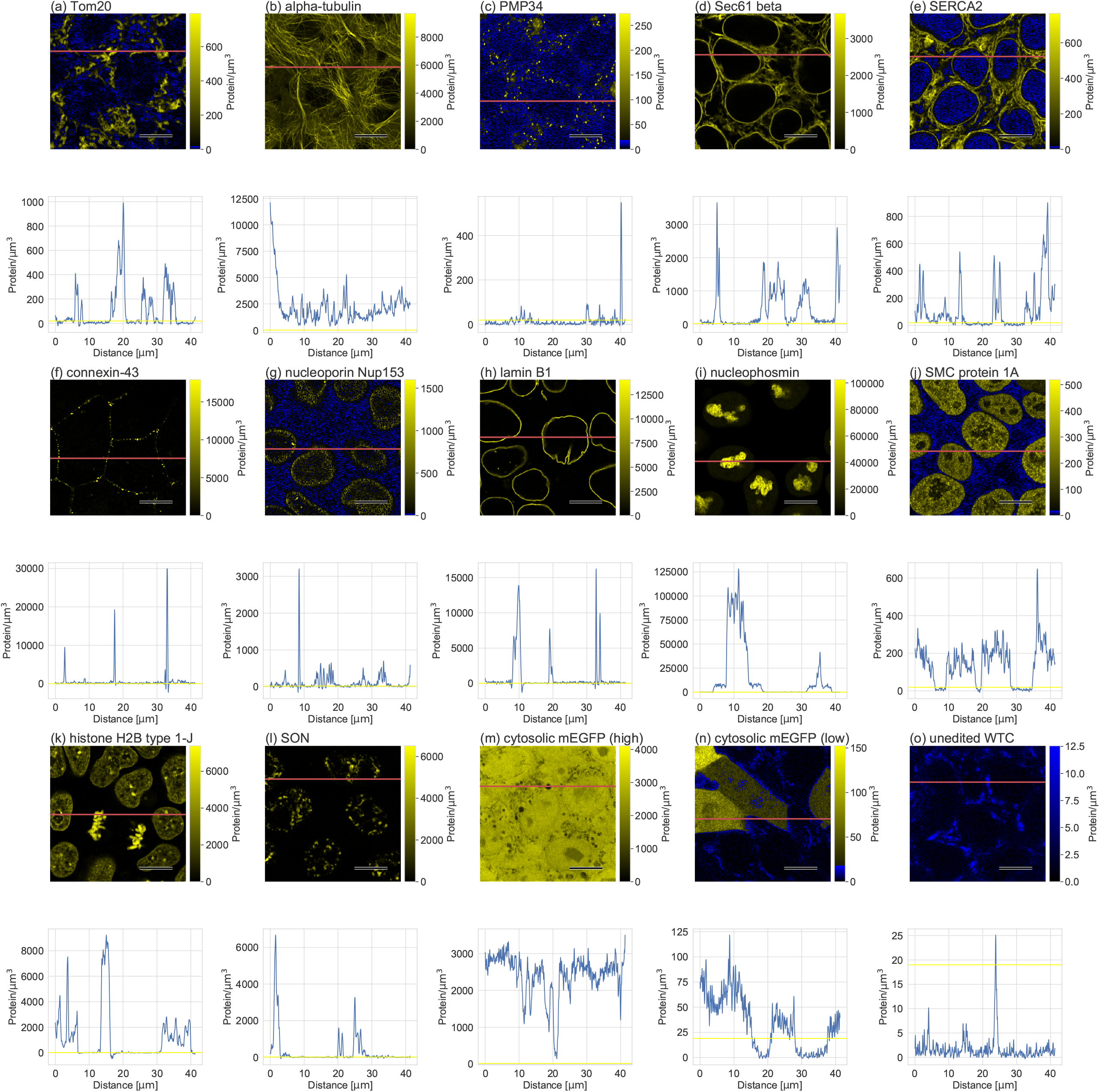
Example of protein number maps for a set of proteins expressed in hiPSCs. All data were acquired as 3D stacks with a laser scanning microscope in Airy Fast mode (only one representative slice per structure is shown, see Supplemental Movies S1-15 for examples of 3D stacks). Protein maps shown represent a variety of key organelles and structures which were endogenously tagged in hiPSC as part of the Allen Cell Collection. The cytosolic mEGFP (high) cell line is the same cell line shown in Figure 1 and Figure 2a. The cytosolic mEGFP (low) cell line is a non-clonal hiPS cell line expressing mEGFP driven by a PGK promoter from the AAVS1 safe harbor locus. The scale bars in the images are 10 μm. The color bars are calibrated as the total number of proteins per μm^3^ and show a different range for each protein. All values in blue are below the level of detection (LOD). The red lines indicate the position of the protein number profile shown below each image. Profile: Each profile was collected along the line in red in the protein number map above. The yellow line indicates the LOD.

### Validation using cells expressing cytosolic mEGFP

We used hiPSC lines expressing free, cytosolic mEGFP to compare the mean cytoplasmic protein concentration determined by our image-based method with results from point-FCS and immunoblotting (Figure 2a). We acquired 3D image stacks with an LSM. After calibration, we calculated the molecular number from all pixels above the background, which was determined from z-slices outside of cells. This simple approach afforded an intrinsic cellular segmentation because the mEGFP signal inside the cells was more than 100-fold above background, resulting in an average concentration of 2.1 ± 0.2 μM or 1,300 ± 100 molecules/μm^3^ for mEGFP in the hiPSC line used for the measurement. We compared this estimate to concentrations determined by point-FCS from 12 cells (one position/cell) in cytosolic regions with uniform fluorescence intensity. The average concentration from these measurements was 1.9 ± 0.6 μM or 1,100 ± 400 molecules/μm^3^. Finally, we compared these results to four separate estimates using quantitative immunoblots. To determine the mean concentration per cell, we calculated the amount of protein per cell divided by the average cell volume (1,700 ± 45 μm^3^ obtained from estimates using 125,000 segmented cells [19] yielding a value of 1.6 ± 0.1 μM or 960 ± 60 molecules/μm^3^).

### Correction for mono-allelic mEGFP labeling

Most of the cell lines used for this study expressed a monoallelic mEGFP tag. As part of the quality control process when generating these cell lines, the relative amounts of untagged vs. tagged protein are assessed by immunoblotting whole cell extracts using protein-specific antibodies ([13] and see Cell Catalog section on www.allencell.org). We then corrected the protein number determined for mEGFP by this factor to calculate the total protein numbers per μm^3^ (mEGFP-tagged + untagged protein) (Supplemental Table S1).

In general, the ratios of tagged protein to total protein (labeling ratios) were close to 0.50 for most of the proteins we imaged. However, for some proteins, the tagged allele was expressed at lower levels than the untagged allele. For example, fibrillarin had a labeling ratio of 0.23 and nucleophosmin had a labeling ratio of 0.22 ([13] and see Cell Catalog section on www.allencell.org). This suggests an adaptive response of cells to a potential functional perturbation of the tag [13].

In addition to differential expression, we also determined whether the ratio of tagged to untagged proteins incorporated into various structures is in proportion to their relative expression levels. Using a multi-step fractionation method (general and organelle specific, see Methods), we blotted all three fractions and determined the molar ratio between the tagged and untagged protein across the different fractions. For the six proteins we tested, we did not see a significant difference in the ratios across fractions suggesting that tagged proteins were not excluded from its structure (Supplemental Table S2 and Supplemental Figure S3).

### Mapping protein numbers

We used this method to create 3D maps of protein numbers using several cell lines from the Allen Cell Collection (www.allencell.org) that express endogenously tagged proteins (Figure 3 shows representative single z-slices, full 3D protein maps can be found in Supplemental Movies S1-15). For regions of the cytoplasm or structures that cannot resolve optically, we express the maps as protein number per μm^3^. For fluorescent proteins residing in supramolecular structures, the numbers are expressed as protein number per structure. While some structures might contain cytoplasmic/nucleoplasmic contributions in the imaged volume element, in general, this latter contribution is low for structures used in this study. For example, SMC protein 1A, part of the nuclear cohesin complex (Figure 3j) had granular appearance throughout the nucleus with numbers above 180 proteins/μm^3^ without any detectable signal outside the nucleus. A manual segmentation of the nucleus allowed reliable integration of the signal, producing a total average protein estimate of 94,000 ± 14,000 molecules in the nucleus. Our estimate was similar to the estimated abundance of another cohesin subunit Rad21 (109,400 molecules per nucleus) in mouse ES cells [20].

Our 3D protein number map allows quantifying the number of proteins localized in distinct subcellular compartments. As expected, membrane anchored proteins tend to have very low protein numbers in the cytosol. The cytosolic density of Tom20, for example is less than 5 molecules/µm^3^ (Figure 3a), which is below the LOD. The numbers of nuclear proteins (e.g., histone H2B type 1-J, SMC protein 1A) in the cytoplasm are ∼ 10 molecules/µm^3^ (Figure 3j, k), consistent with the presence of its nuclear localization sequence that concentrates the protein in the nucleus [21]. In contrast, the density of cytosolic alpha-tubulin is ∼ 1,000 molecules/µm^3^ (Figure 3b), indicating a substantial amount of soluble tubulin for maintaining microtubule assembly and disassembly [22]. Some proteins that are enriched in membrane-less nuclear substructures also have a substantial pool in the nucleoplasm. For example, ∼ 7,000 molecules/µm^3^ of nucleophosmin are present in the nucleoplasm with ∼ 100,000 molecules/µm^3^ in the nucleolus (Figure 3i). Thus, nucleophosmin concentration in the nucleoplasm is approximately 14-times lower than in the nucleolus, but substantially higher than other nucleoplasmic proteins we analyzed such as SMC protein 1A (Figure 3j). The large dynamic range of local protein densities in living cells should allow quantitative analyses and computational models for subcellular compartmentalization and its regulation in the context of nucleocytoplasmic transport and intracellular phase separations, for example.

From these findings, we conclude that our method provides estimates of protein numbers in several cellular structures as well as the numbers of proteins per cell, facilitating comparison of protein abundance in different subcellular compartments. These maps will become particularly useful when combined with numbers of other proteins to determine how they might pack in complex structures.

### Extension to other microscope modalities

To generalize the method, we adapted it to two other commonly used microscope modalities in addition to LSM (Figure 1, Figure 2a). SD microscopes offer high imaging throughput and robustness, and the LLS microscope is better for dynamic image information due to its high acquisition rate and low phototoxicity.

In contrast to the LSM, both the SD and LLS have uneven illumination (Figure 2d, f). To address this, we created a calibration curve for each xy pixel in the image plane of SD and LLS data to remove uneven illumination artifacts (Figure 2d-g). Because of its multiple pinholes, the SD collected out-of-focus light when imaging >200 μm above the cover slip. For the SD, we quantified the contribution of the pinhole crosstalk (Figure 2c). At 200 µm above the coverslip in SD, fluorescence intensity of EGFP solution changes minimally with changes in the z-position (Figure 2c). Therefore, we acquired calibration images at 200 µm above the coverslip and corrected for pinhole crosstalk in the calibration curve accordingly. On LSM and LLS microscopes, fluorescence intensities of LSM and LLS images were constant along the z-axis (Figure 2c, data not shown). For consistency, we chose 200 µm above the coverslip to acquire EGFP calibration images on LSM and LLS microscopes as well. After we applied the microscope-specific calibration procedures discussed above to cells expressing cytosolic mEGFP, all instrument specific differences in the images disappeared and only biological variances remained (Figure 1c-e), showing that the method can be used across microscope modalities when appropriate adjustments are made.

We applied the methods used to determine detection limits for LSM to the SD and LLS microscopes (Supplemental Figure S4). While the LSM could potentially measure concentrations > 10-fold lower than SD and LLS platforms, the autofluorescence made differences between LSM and SD negligible. LLS performance was the least affected by autofluorescence and scattering. The parallel detection of SD microscopes that resulted in pinhole crosstalk for thick samples, as discussed above, resulted in the benefit of a higher precision compared to the higher resolution of LSM.

## Summary

We implemented and validated a robust, scalable, and easy-to-use method for calibrating fluorescence intensities to measure the numbers of mEGFP-tagged proteins in living cells from 3D images across three different microscope modalities. We used hiPSC lines in which proteins of interest were endogenously tagged (monoallelic) with mEGFP. Thus, the measured fluorescence intensity of mEGFP was proportional to the number of tagged proteins. By correcting for the ratio of tagged to total protein, we quantified the total (untagged + tagged) numbers for the protein of interest. A major advantage of the presented method was that it generated 3D maps of absolute protein number in single cells at imaging resolutions suitable for subcellular compartments. Compared with previous studies in mapping protein number in cells, this method has introduced two key developments of live cell imaging: 1) enhanced spatial resolution by LSM Airyscan imaging; 2) long term 3D imaging by LLS. As we have demonstrated, this approach reliably measures protein number in multiple cell lines with protein densities across three orders of magnitudes. The robust calibration algorithm enabled handling large volume of image data and generated protein number maps within a short processing time (usually less than 20 minutes for 50 image z-stacks). A similar approach could be used in dual or triple edited cells to create multidimensional maps of protein number.

The protein number data enable answers to a multitude of questions including: What are the magnitudes and origins of the cell to cell variation in protein expression? What is the relation between total protein number and cell size and how does it vary as cells change behavior? What is the relation between gene expression and protein number in single cells? What is the fraction of bound vs soluble protein, e.g., the equilibrium constant for protein associations with components of the cellular machinery? All these potential applications are driven by the ability to easily measure absolute, local numbers of proteins with sub-cellular resolution in a large, statistically meaningful number of single cells, using microscope modalities optimized for specific tasks, and to make meaningful statistical interferences using this image data.

## Acknowledgements

This work was only possible with the help of many individuals from the Allen Institute for Cell Science who created and characterized the cell lines, provided and maintained the imaging infrastructure, and helped with data dissemination and project management. We thank Julie Theriot, David Piston, and Enrico Gratton for helpful discussions. The parental unedited hiPSC cell line, WTC, was provided by the Bruce R. Conklin Laboratory at the Gladstone Institutes and UCSF. We wish to thank the Allen Institute founder, Paul G. Allen, for his vision, encouragement and support.

## Material and Methods

### Gene-edited cell lines

All clonal human induced pluripotent stem cell (hiPSC) lines in this study are from the Allen Cell Collection (www.allencell.org) and are listed with brief descriptions of the mEGFP labeled proteins in Supplemental Table S1. The cell lines were cultured on Matrigel (Corning, #356231) in mTeSR1 media (Stemcell Technologies, cat #85850) and seeded into Matrigel-coated 96-well glass bottom imaging plates (Cellvis, #P96-1.5H-N) through a Microlab STAR liquid handling system (Hamilton). Cells were imaged in phenol-red free media (Stemcell Technologies, # 05876) at 37°C and 5% CO_2_ [23].

### EGFP stock solution and dilution series

We used *E. coli* purified EGFP (Chromotek, #EGFP-250) as a stock solution. The stock concentration of 1 ± 0.1 g/L = 35 μM as specified by the vendor was confirmed by immunoblotting as well as by measuring the absorbance at 280 and 488 nm using a Nanodrop 8000 Spectrometer (ThermoFisher Scientific). We used the molar absorptivity coefficients of 22,015 L/(mol · cm) (based on amino acid composition) for 280 nm and 55,000 L/(mol · cm) for 488 nm [24] to calculate the concentration of the stock solution from the absorbance measurements.

The stock solution was diluted in 1:20 in PBS (pH 7.4), 0.1% BSA, 14 mM NaN_3_ and 1:500 ProClin 300 (Millipore Sigma, #48912-U) to create a working stock of 1.75 μM. The working stock was serially diluted to final concentrations of 875, 583, 194, 65, 22, 7 nM, including a blank in a 96-well glass bottom imaging plate. The EGFP concentrations of the standard curve were measured with FCS in parallel to each experimental imaging session of hiPSCs (for details see “FCS in solution”). We used FCS to determine the concentrations because this method can measure the concentration range we are interested in (Supplemental Figure S2) and could be used with the small sample volume from our dilution series. Therefore, we could confirm the concentrations daily and use the same dilution series up to a few weeks without having to worry about changing concentrations caused by buffer evaporation or protein degradation and adsorption on container surfaces.

### Microscopy

LSM data were acquired in the “fast” mode using the Carl Zeiss LSM 880 Airy Fast inverted confocal laser scanning microscope [25] with a C-Apochromat 40x/1.2NA water-immersion objective, 884 × 884 pixels (0.05 × 0.05 μm^2^) and 65 z-slices (0.2 μm spacing). EGFP was excited with a 488 nm Argon-ion laser at 0.5-4% power level (equal to 11-77 μW) measured with the C-Apochromat 40x/1.2NA water-immersion objective. The detector was a 32-channel gallium arsenide phosphide (GaAsP) Airy detector operated in the 16-channel fast mode with pixel dwell times 2.45-4.9 μs and gains 850-1,000 dependent on observed intensity levels. Unless noted otherwise, all data were processed in the Zeiss ZEN Black software using the 3D Airy deconvolution with the filter strength selected automatically. Images of the EGFP dilution series were acquired 200 μm above the coverslip.

For SD microscopy, we used a CSU-X1 scan head with 50 μm pinholes (Yokogawa) integrated in an inverted imaging platform (Carl Zeiss, Cell Observer SD). All data were acquired with a C-Apochromat 100x/1.25NA water-immersion objective and two Orca-Flash 4.0 v2+ sCMOS cameras (Hamamatsu), 924 × 624 pixels after 2×2 binning (0.11 × 0.11 μm^2^) and 200 ms exposure time. Each 3D image stack had 75 z-slices (0.29 μm spacing). mEGFP was excited at 488 nm with a power of 2.3 mW (measured with a 10x objective) from a diode laser. Transmitted light images were acquired simultaneously with 740 nm LED illumination. Images of EGFP dilution series were acquired 200 μm above the coverslip.

An LLS microscope (Carl Zeiss, Lattice Lightsheet 7) was equipped with a custom optical illumination (13.3x/0.44NA) and detection (44.83x/1.0NA) unit to generate a 30 μm × 1 μm light-sheet that enters the sample at an angle of about 30° relative to the cover slip. Samples were excited with a solid-state 488 nm diode laser (160 μW laser power) and images were collected with an sCMOS camera (PCO, PCO-Edge) with an exposure time of 100 ms per frame. 501 sections were acquired by moving the stage along the y-axis in 0.2 μm steps. Pixel size was 0.145 × 0. 145 μm^2^ without binning. Image stacks were de-skewed using the “coverglass transformation” option in ZEN Blue 3.1 acquisition software before calibration analysis.

### FCS in solution

All FCS data were acquired with a Zeiss LSM 880 Airy fast confocal microscope with a C-Apochromat 40x/1.2NA water immersion objective with manual or motorized correction collar (Carl Zeiss). EGFP in solution was excited with 488 nm light from an Argon-ion laser (laser power 1.8 μW) and detected through a bandpass filter (BP 495-621 nm) with a GaAsP detector in single photon counting mode (count rate 20 MHz) (see Supplemental Figure S2a for photon traces). We measured each concentration three times for 180 s. The pinhole was set to one Airy unit and the correction collar was optimized for maximum counts-per-molecule or minimum width of reflection profile from the cover slip along the optical axis. The intensity signal was autocorrelated using the software auto-correlator of the FCS module for the Zeiss ZEN Black software (see example in Supplemental Figure S2b). Least square fitting of FCS models to autocorrelated signal was performed with ZEN Black or the Python package LMFIT [26]. Our autocorrelation profiles revealed at least two distinct components. Previous studies suggest that fluorophore blinking is the source of the faster components while the diffusion of the fluorophore dominated the slower [27, 28]. Therefore, we explored multiple fitting models and strategies. All models were of the form:

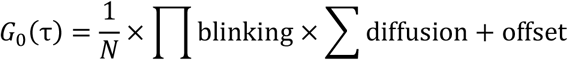

With blinking terms in the form

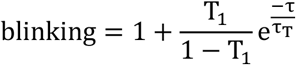

diffusion terms of the form

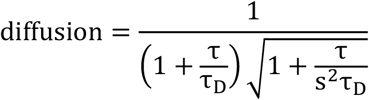

and a constant offset. The correlation time τ was the variable and the fit was used to determine number of molecules N, triplet (blinking) fraction T, triplet (blinking) time constant τ_T_, diffusion time constant τ_D_, and offset o. The structure parameter s was set to a fixed value of 10.

To process FCS data for calibration curves, we used a model with two blinking and one diffusion component, as it worked best with our data.

We used intensity recordings from buffer solution without EGFP (blank buffer) to correct N for background signal [27]:

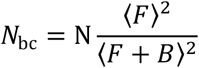

with mean fluorescence intensity ⟨F⟩ and mean sum of fluorescence and background signal ⟨F + B⟩. We calculated EGFP concentration c according to

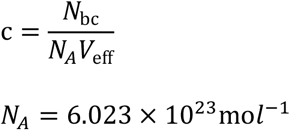

with effective volume V_eff_ for short and long axis ω_xy_ and ω_z_ [29, 30]:

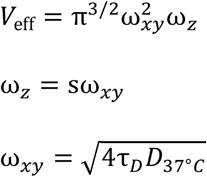

We calculated EGFP number per μm^3^ according to

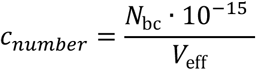

when *V*_*eff*_ is expressed in the unit of liter. We used *C* for generating calibration curves shown in Figure 2 and *C*_*number*_ for generating calibrated images in Figure 1 and 3.

We corrected the diffusion constant for EGFP at 22.5°C of D_22.5°C_ = 95 μm^2^/s [31] for 37°C [32]:

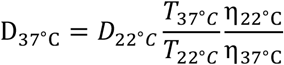

The viscosities for water at 22.5°C was η_22.5°C_ = 9.43×10^−4^*Pa*·s and at 37°C it was η_38°C_ = 6.91×10^−4^*Pa*·s (retrieved from WolframAlpha and based on National Institute of Standards and Technology, NIST Reference Fluid Thermodynamic and Transport Properties Database (REFPROP)) and T the temperatures in Kelvin.

### FCS in hiPSC

We used an LSM 880 with GaAsP detector (Carl Zeiss) in single photon counting mode to measure concentrations of mEGFP in the cytosol of hiPSC using FCS. We imaged cells in confocal imaging mode and positioned the laser beam in the cytosol of cells in areas of even fluorescence to avoid visible organelles. At these positions, we acquired one point-FCS curve per cell and excluding all data showing motion or bleaching artifacts. These artifacts indicated that the cell or subcellular structures had moved and thus the data was not reliable. mEGFP was excited with 488 nm at 0.2% laser power setting and AOTF damping of 7.5 through a C-Apochromat 40x/1.2NA water immersion objective with manual correction collar. Data was acquired for 30 s per cell and correlated using ZEN Black software. We fitted a one component blinking and diffusion to the autocorrelated data between 1 µs and 30 s correlation times. We calculated the local concentration using the approach described for FCS in EGFP solution.

### Concentration calibration of fluorescence images

We used our EGFP dilution series to calibrate our microscopes. First, we optimized the microscope settings (e.g., objective, laser power, detector gain, exposure time) to image mEGFP-labeled structures in hiPSCs. Then, we used the exact same settings to image the EGFP calibration curve (preparation described above). We used a linear regression to create the intensity – concentration calibration curve.

From the concentrations c_Well_ determined by FCS for the EGFP dilution series, we subtracted the offset given by the apparent concentration for the blank buffer well c_Buffer_:

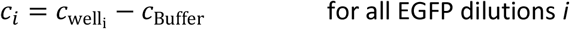

From LSM images of EGFP dilution series we excluded 5% of edge pixels and calculated the average ⟨⟩ over the whole image. We subtracted the average intensity o of the buffer blank well from the images of EGFP buffer dilution series:

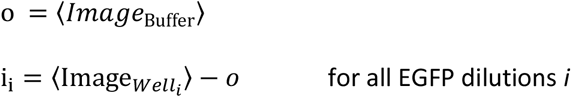

The calibration factor a was calculated as

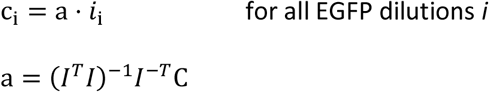

With I and C vectors of all i_i_ and c_i_ and T the matrix transforms.

For LSM images I_LSM_ we subtracted the mean of the blank buffer well o from each pixel of the original image I_LSM_ and multiplied it by the calibration factor a:

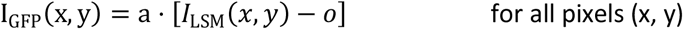

to get an image calibrated in mEGFP concentrations I_GFP_.

Finally, we divided the 3D stack of I_GFP_(x, y) by the labeling ratio for each protein to obtain the 3D stack of total protein counts.

Although we used concentrations determined by FCS for the EGFP dilution series, concentrations measured by other methods (such as absorption spectroscopy) can be used instead in the algorithm to generate calibration curves and protein count maps. In these cases, EGFP stock concentration can be measured by spectroscopy and an accurate dilution series needs to be prepared in order to generate a good calibration curve.

To take instrument-specific noise sources into account, we imaged buffer solutions without any dye and determined the 99% confidence interval of the background signal (Supplemental Figure S4a), which was equivalent to 3.8 proteins per μm^3^ (99% of all background counts are below this value). We multiplied this value by five (a value typically used to establish LOD) and established 19 proteins/μm^3^ as the LOD.

Because of the uneven illumination in SD and LLS we had to adapt the calibration procedure. We could no longer use the same offset and slope for the whole image and therefore calculated them for each pixel in the image plane individually (Figure 2d-g).

It is known that fluorescence quenching of EGFP was significant upon changes in solution pH [28]. Although the lumen of some organelles such as lysosomes and the Golgi apparatus has lower pH [33] that could cause EGFP quenching, all proteins we have examined have the fused mEGFP located to the cytoplasmic side of organelles, in the cytoplasm or the nucleoplasm. Therefore, our results are not subject to EGFP quenching caused by acidic pH. Because most of our tagged proteins are in 0.1-10 µM range, we consider the effects of protein crowding on our concentration measurement to be negligible. While these estimates could be influenced by molecular crowding, the effect is likely small since EGFP homotetramers showed only a 4% decrease in fluorescence lifetime when compared to GFP monomers [34].

### Organelle fractionation and immunoblotting

Cells were dissociated using Accutase (Stemcell Technologies, #07920) following the standard procedures [13], counted using a Vi-Cell cell counter (Beckman Coulter) and centrifuged for 5 min at 300 × *g*. The cell pellet was resuspended in 200 μL M-PER mammalian protein extraction reagent (ThermoFisher Scientific, #78501) at 50,000 cells per μL lysis solution plus 1x HALT protease inhibitor mix (ThermoFisher Scientific, #78430) and 250 U/ml nuclease (ThermoFisher Scientific, #88702) with 2 mM MgCl_2_, lysed for 10 min at 4°C followed by centrifugation at 16,000 × *g* for 10 min at 4°C. The supernatant was saved for immunoblotting. The insoluble pellet fraction was resuspended in 200 μL Mem-PER Plus membrane protein buffer (ThermoFisher Scientific, #89842) supplemented with HALT protease inhibitor followed by 10 min centrifugation at 16,000 × *g* at 4°C. The supernatant was saved for immunoblotting. Finally, the remaining insoluble pellet fraction was resuspended in 200 μL of 8 M urea (Invitrogen, #15505-035) 2x LI-COR loading buffer solution. Intercept (PBS) protein-free blocking buffer supplemented with 0.2% tween-20 was used for all blocking and antibody staining steps (LI-COR, #927-70001). For microtubules specific fractionation we used a microtubules/tubulin *in vivo* assay kit (Cytoskeleton, #BK038) according to vendor specifications. To enrich nuclear proteins, we used a minute cytoplasmic & nuclear extraction kit for cells (Invent Biotechnologies, #SC-003) according to vendor specifications.

The three lysate fractions were electrophoresed on NuPAGE™ 4 to 12%, Bis-Tris 17-well gels (ThermoFisher Scientific, #NP0349BOX) and transferred to 0.2 μm pore sized nitrocellulose membranes (ThermoFisher Scientific, #LC2000). Odyssey chameleon duo molecular weight markers (LI-COR Biosciences, #928-60000) were used as a molecular size standard. Membranes were incubated overnight at 4°C with anti-protein specific antibodies (see Supplemental Table S2) followed by incubation for 1 hour at room temperature with goat anti-mouse or goat anti-rabbit IRDye 800 (LI-COR, #926-32210, 926-32211) secondary antibodies diluted at 1:5,000. The immunoblots were imaged with an Odyssey CLx imaging system and analyzed with Image Studio Lite 5.2.5 software version (both LI-COR). We report the mole fraction of mEGFP tagged protein over all detected protein bands with mEGFP-tagged and untagged protein.

### Immunoblotting to determine intracellular mEGFP concentration

hiPSCs expressing cytosolic mEGFP (Figure 3m) were lysed using 50,000 cells per μL M-PER mammalian protein extraction reagent as described in the section above. The cell lysate was diluted 2.5-fold into 1x LI-COR loading buffer (LI-COR, # 928-40004) and immunoblots were performed as described above. We used an anti-EGFP primary antibody diluted at 1:1,000 (Sigma, #11814460001) for staining the blot and imaged with Odyssey CLx imaging system.

An EGFP standard curve was obtained by diluting the 1,000 ng/μL EGFP stock solution 1:250 for the top standard curve point in LI-COR 1x loading buffer diluent mix. All subsequent points on standard curve are 3-fold dilutions of the top standard. Final amounts of 40.00, 13.33, 4.44, 1.48, 0.49, and 0.16 ng were run on the same gels as the cell lysates for immunoblotting. We used a linear fit to the standards and integrated band intensities to determine the moles of EGFP per μL cell lysate. We used the number of cells per μL of cell lysate and the median cell volume of 2 pL to calculate the average concentration of mEGFP per cell.

### Access to materials and data

All clonal cell lines used in this study are described in detail in the Cell Catalog section on www.allencell.org and can be obtained through the Coriell Institute at www.coriell.org/1/AllenCellCollection for scientific use. Donor plasmids used to create these gene-edited hiPSC lines can be obtained through Addgene at www.addgene.org/The_Allen_Institute_for_Cell_Science. Data used to produce figures in the paper and Jupyter notebooks for data analysis are available upon request.

