## Supplemental Material for "Mapping Protein Numbers in Living Cells"

**Supplemental information in this PDF includes:**

Table S1-S2
Figures S1 – S4
Captions for Movies S1 - S15

**Other Supplementary Materials for this manuscript include:**
Movies S1 – S15

### Supplemental Tables

| Protein | Gene | Structure | Tag Location | Tagged Alleles | Labeling Ratio |
|---|---|---|---|---|---|
| **SERCA2** | ATP2A2 | Sarcoplasmic reticulum/endoplasmic reticulum | N-terminus | monoallelic | 0.48 |
| **Connexin-43** | GJA1 | Gap junctions | C-terminus | monoallelic | 0.44 |
| **Histone H2B type 1-J** | HIST1H2BJ | Histones | C-terminus | monoallelic | Not measurable, use ratio of 0.5 |
| **Lamin B1** | LMNB1 | Nuclear envelope | N-terminus | monoallelic | 0.33 |
| **Nucleophosmin** | NPM1 | Nucleolus (granular component) | C-terminus | monoallelic | 0.22 |
| **Nucleoporin Nup153** | NUP153 | Nuclear pores | N-terminus | monoallelic | 0.47 |
| **Sec61 beta** | SEC61B | Endoplasmic reticulum | N-terminus | monoallelic | 0.53 |
| **Peroxisomal membrane protein PMP34** | SLC25A17 | Peroxisomes | C-terminus | monoallelic | 0.49 |
| **SMC protein 1A** | SMC1A | Cohesins | C-terminus | monoallelic (on X chromosome in male iPSC background so only one gene copy) | 1.0 |
| **SON** | SON | Nuclear speckles | N-terminus | monoallelic | Not measurable, use ratio of 0.5 |
| **Tom20** | TOMM20 | Mitochondria | C-terminus | monoallelic | 0.54 |
| **Alpha-tubulin** | TUBA1B | Microtubules | N-terminus | monoallelic | 0.55 |

**Table S1. Protein names, gene names and corresponding structures, for all proteins used in this study.** "Tag location" describes the protein terminus tagged with mEGFP and "Tagged alleles" the allelic count of the tag. The labeling ratio is the ratio between mEGFP-tagged proteins and total proteins as determined by immuno-blot information from (13) and cell catalog on www.allencell.org.

| Protein | Gene | GFP-protein Mw [kDa] | Method | Fraction 1 | Fraction 2 | Fraction 3 | Fraction 2 + 3 | Vendor | Antibody | Dilution |
|---|---|---|---|---|---|---|---|---|---|---|
| Tom20 | TOMM20 | 47 | General | 22+/-5 | 23+/-5 | 9+/-3 | 19+/-5 | Santa Cruz Biotechnologies | sc-17764 | 1:250 |
| Alpha-tubulin | TUBA1B | 77 | Cytoskeleton | 40+/-3 | 42+/-4 | | | ThermoFisher | 62204 | 1:10000 |
| Nucleoporin Nup153 | NUP153 | 180 | Nucleus | 56+/-20 | 64+/-20 | | | Sigma-Aldrich | HPA027897 | 1:250 |
| Lamin B1 | LMNB1 | 93 | Nucleus | 47+/-2 | 45+/-2 | | | Abcam | ab16048 | 1:2000 |
| Sec61 beta | SEC61B | 37 | General | 68+/-3 | 96+/-1 | 57+/-8 | 61+/-7 | Abcam | ab15576 | 1:10000 |
| SON | SON | 291 | Nucleus | 54+/-10 | 88+/-2.7 | | | Sigma-Aldrich | HPA023535 | 1:500 |

**Table S2. Proteins used for level of incorporation measurements.** The type of incorporation measurements is listed as general for the organelle unspecific fractionation method or with the organelle name for organelle specific methods. The three fractions are listed for the general method (Fraction 1: M-PER cytoplasmic fraction; Fraction 2: Mem-PER Plus membrane protein enriched fraction, Fraction 3: urea highly insoluble protein fraction) and two fractions are listed for the organelle specific fractionation kits (Fraction 1: non-organelle or non-polymerized fraction; fraction 2: organelle or polymerized fraction). The primary antibody vendors, product numbers, and dilution factors are described in the table.

### Supplemental Figures

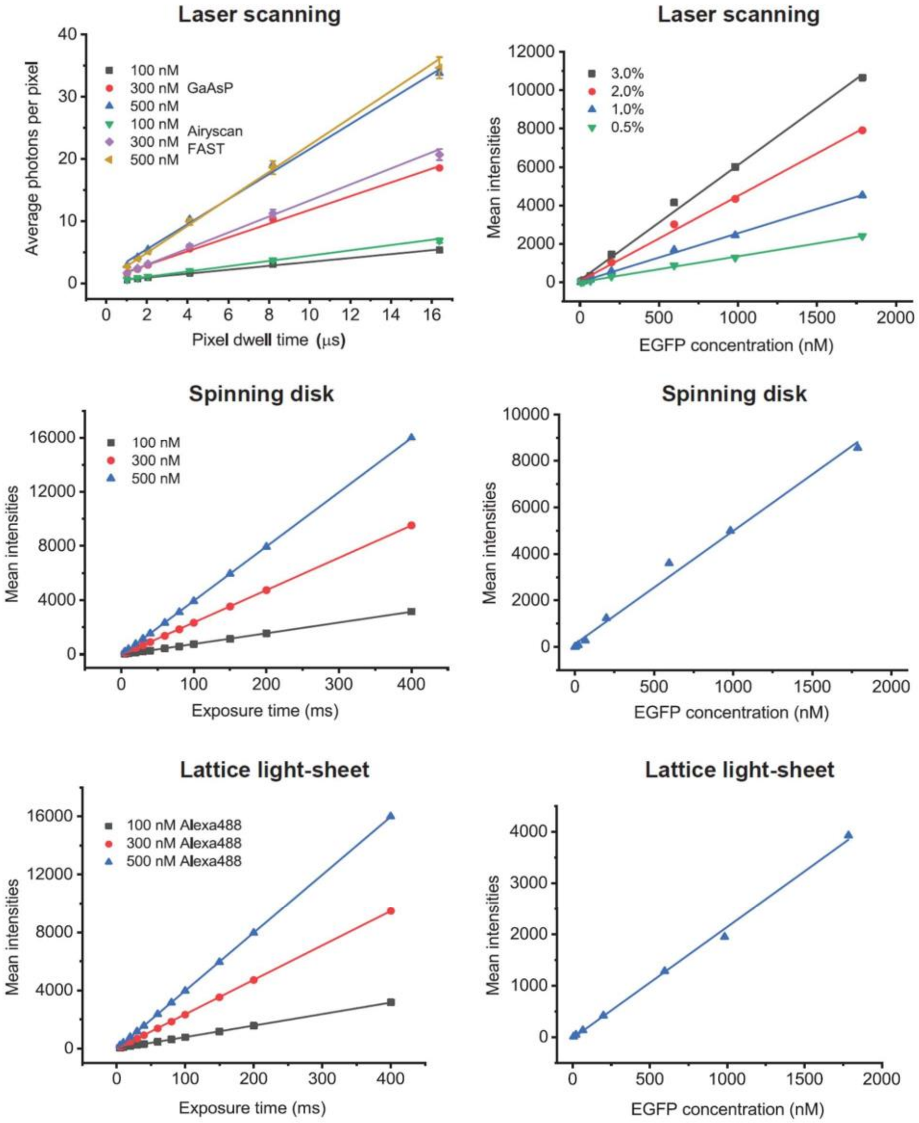


**Figure S1. Linear detector response for different exposure times or different EGFP concentrations** (Left column) Alexa-488 dye solution was imaged with different pixel dwell times or camera exposure times, while keeping all other imaging parameters constant. Increasing pixel dwell times or camera exposure times results in a proportional increase of fluorescence signal at each microscope detector. Linear fits to the data are shown. (Top) Photon counts of imaging Alexa Fluor 488 at 100 nM, 300 nM and 500 nM concentrations from a single channel GaAsP detector (used for FCS in this work) or in Airy Fast mode (used for EGFP calibration imaging) with a laser scanning microscope. Pixel dwell time range from 1.02 µs to 16.38 µs. The single channel GaAsP detector was operated at pinhole size of 0.2 Airy Unit. The average of 8 central detectors in Airy FAST mode was plotted against pixel dwell times. In all cases the $R^2$ value for the linear fit (lines) was higher than 0.996. Excitation with 61.8 µW laser power at ChA FAST is equivalent to excitation with 20.7 µW laser power at GaAsP. (Middle and bottom). Mean intensity on an sCMOS camera of confocal spinning disk microscope (middle) and on an sCMOS camera of lattice light-sheet microscope (bottom) for Alexa 488 dye solutions with concentrations of 100 nM (black squares), 300 nM (red circles), and 500 nM (blue triangles). Exposure times range from 5 ms to 400 ms. In all three case the $R^2$ value for the linear fit (lines) was higher than 0.999. (Right column) Mean intensities of EGFP dilution series measured with a laser scanning microscope in Airy Fast mode (top), spinning disk (middle) and lattice light-sheet (bottom). Laser powers used on laser scanning microscope are shown in the plot. Acquisition settings for spinning disk and lattice light-sheet are described in Materials & Methods section. Linear fits for each plot are shown as lines.

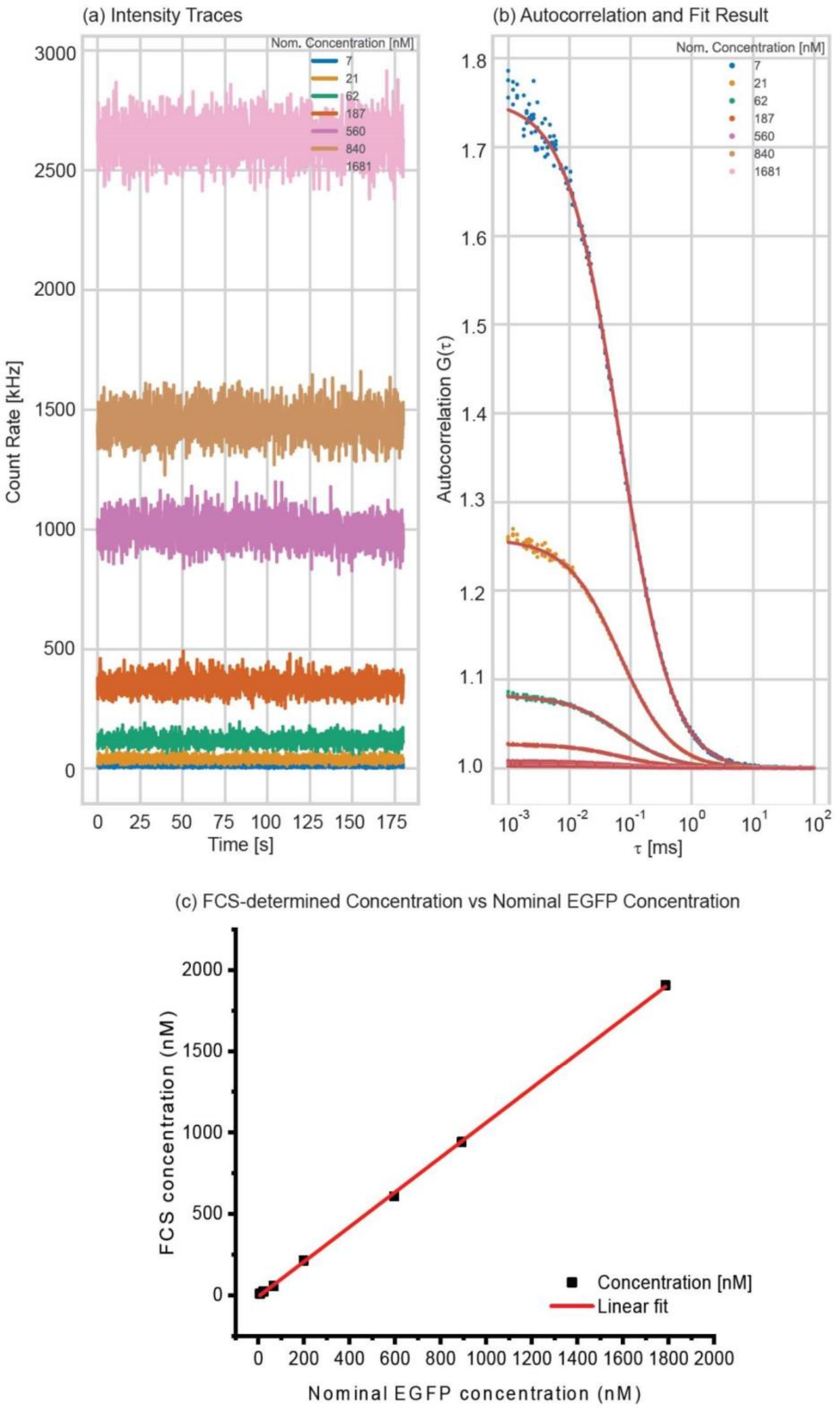


**Figure S2. Calibrating EGFP concentrations by FCS**. **(a)** Traces of single photon count rates acquired with a laser scanning microscope in EGFP solutions of different concentrations (for visualization purposes only a subset of datapoints is shown). **(b)** Autocorrelation G(τ) curves (dots) of fluctuation data shown in (a). Red lines show fit results for one component diffusion model with blinking components (see Methods). Data beyond τ = 100 ms are not shown. **(c)** Plot of FCS-measured concentrations versus nominal EGFP concentrations on one experiment day. The slope of the linear fit is 1.066 ± 0.008, with an $R^2 = 0.9997$.

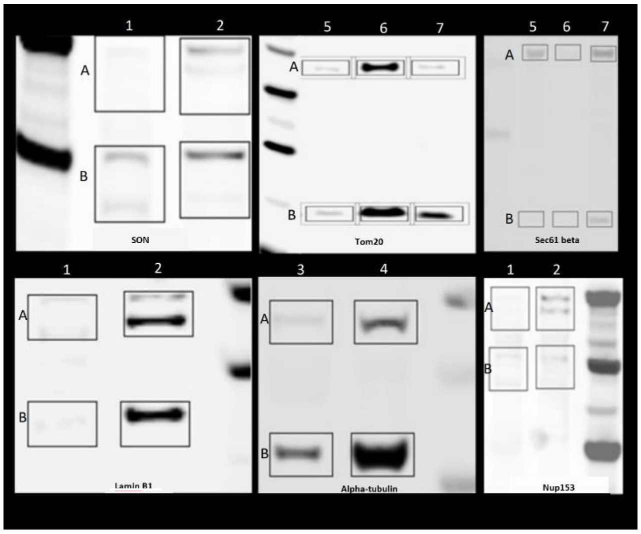


**Figure S3: Immunoblots used to determine behavior of tagged and untagged proteins during fractionations**. Band A is mEGFP tagged protein and band B is untagged protein. Lane: 1, 3 non-organelle or non-polymerized fraction from organelle-specific kits, respectively. Lane 2: nuclear fraction from organelle-specific kit. Lane 4: the polymerized alpha-tubulin fraction form organelle-specific kit. Lane 5: M-PER extraction fraction. Lane 6: Mem-PER Plus fraction. Lane 7: urea fraction.

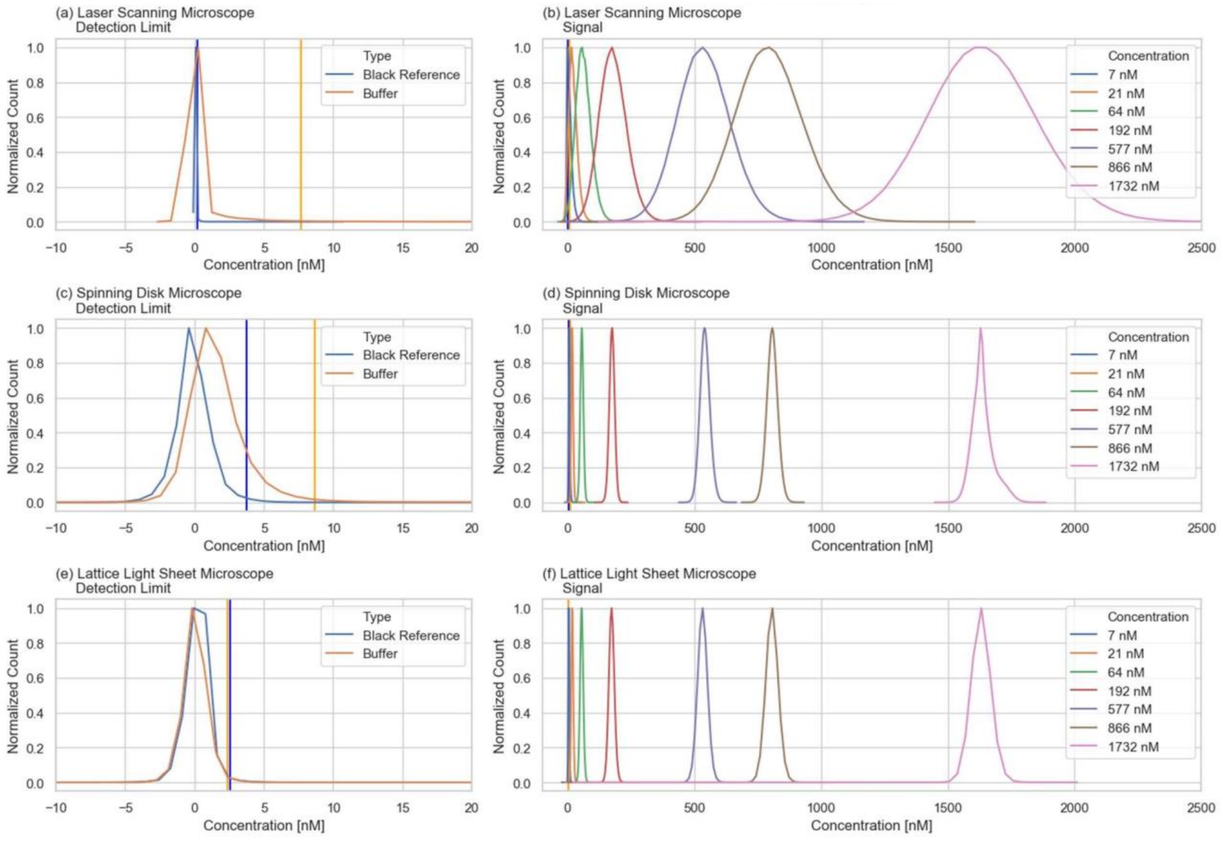


**Figure S4. Detection limits of different microscope systems**. **(a, c, e)** Histograms of concentrations measured without any laser exposure (Black Reference) and buffer solution without any dye (Buffer) using (a) a laser scanning microscope, (c) spinning disk microscope, and (e) a lattice light-sheet microscope. The blue and orange vertical lines indicate the 99% confidence interval (99% of all concentration values are below this value) for black reference and buffer respectively. **(b, d, f)** Histograms for concentrations measured in EGFP solutions with different nominal concentrations using (b) a laser scanning microscope, (d) spinning disk microscope, and (f) lattice light-sheet microscope.

### Supplemental Movie Captions

**Movie S1-S15. 3D z-stacks of protein number maps (in molecules/μm$^3$) of 14 gene-edited hiPSC lines from the Allen Cell Collection and the unedited wild type cell (WTC) line.** Raw images were acquired from a Zeiss LSM880 confocal microscope in Airyscan Fast mode. Movies were compressed and converted to MP4 format using Handbrake 1.3.3. Calibration bars are displayed at the upper right corner of each Movie.

Movie S1. 3D Protein number map of Tom20
Movie S2. 3D Protein number map of alpha-tubulin
Movie S3. 3D Protein number map of peroxisomal membrane protein PMP34
Movie S4. 3D Protein number map of Sec61 beta
Movie S5. 3D Protein number map of SERCA2
Movie S6. 3D Protein number map of connexin-43
Movie S7. 3D Protein number map of nucleoporin Nup153
Movie S8. 3D Protein number map of lamin B1
Movie S9. 3D Protein number map of nucleophosmin
Movie S10. 3D Protein number map of SMC protein 1A
Movie S11. 3D Protein number map of histone H2B type 1-J
Movie S12. 3D Protein number map of SON
Movie S13. 3D Protein number map of cytosolic mEGFP (high)
Movie S14. 3D Protein number map of cytosolic mEGFP (low)
Movie S15. 3D Protein number map of unedited WTC (as background control)
